# Neuronal Subcompartment Classification and Merge Error Correction

**DOI:** 10.1101/2020.04.16.043398

**Authors:** Hanyu Li, Michał Januszewski, Viren Jain, Peter H. Li

## Abstract

Recent advances in 3d electron microscopy are yielding ever larger reconstructions of brain tissue, encompassing thousands of individual neurons interconnected by millions of synapses. Interpreting reconstructions at this scale demands advances in the automated analysis of neuronal morphologies, for example by identifying morphological and functional subcompartments within neurons. We present a method that for the first time uses full 3d input (voxels) to automatically classify reconstructed neuron fragments as axon, dendrite, or somal subcompartments. Based on 3d convolutional neural networks, this method achieves a mean f1-score of 0.972, exceeding the previous state of the art of 0.955. The resulting predictions can support multiple analysis and proofreading applications. In particular, we leverage finely localized subcompartment predictions for automated detection and correction of merge errors in the volume reconstruction, successfully detecting 90.6% of inter-class merge errors with a false positive rate of only 2.7%.

## 1 Introduction

Recent advances in 3d electron microscopy (EM) have enabled synaptic-resolution volumetric imaging of brain tissue at unprecedented scale [1–3]. Semi-automated reconstructions of these volumes yield thousands of neurons and neuronal fragments, inter-connected by millions of synapses [4–7]. Together, reconstructed neurons and synapses within each dataset describe a “connectome”: a connectivity graph whose structure is anticipated to underlie the computational function of the tissue [8, 9].

Interpreting neural connectivity at this scale is a significant undertaking. One means to enhance interpretability is to use ultrastructural and morphological details of neuronal fragments to distinguish their functional subcompartments. For example, the classical description of “neuronal polarity”, i.e. the flow of information within vertebrate neurons, from dendritic subcompartments, into the soma, and out through the axon, remains central to understanding connectivity [10].

Although trained human reviewers can classify many neuronal fragments with respect to subcompartment, the growing scale of connectomic reconstructions demands automated methods. A recent approach to this problem was based on training random forest classifiers on manually defined features extracted from neurite segments and separately detected organelles such as mitochondria or synapses [4, 11]. A later extension improved accuracy by classifying 2d projections of neurites and their organelles with convolutional neural networks (CNNs), a technique called Cellular Morphology Networks (CMNs) [12]. However, an approach based on a full 3d representation of neuron fragments, which retains the maximum morphological and ultrastructural information, has not been previously demonstrated.

Another application for subcompartment predictions is in proofreading, e.g. to correct errors in automated reconstructions. Prior works proposed to detect merge errors through identification of morphologically unlikely cross-shaped fragments [11, 13] or used a 3d CNN trained specifically to detect merge [14] or split errors [15]. Strong biological priors dictate that vertebrate neurons have only one major axonal branch extending from the soma, and that dendritic and axonal subcompartments do not typically intermingle within a neurite [10]. Prior work used these cues to tune agglomeration via sparse subcompartment predictions in a multicut setting, which optimizes over an explicit edge-weighted supervoxel graph [16, 17]. Alternatively, violation of biological priors in subcompartment predictions can be used to detect post-agglomeration reconstruction errors, which can then be flagged for efficient human proof-reading workflows [18], or fully automated error correction.

In the following we (1) present a system for neuronal subcompartment classification based on 3d convolutional neural networks, (2) demonstrate finely localized subcompartment predictions whose accuracy exceeds state-of-the-art, and (3) show how these predictions can be used for high-fidelity detection and correction of agglomeration errors in an automated segmentation.

## 2 Materials & Methods

### 2.1 Datasets

We used an automated Flood-Filling Network (FFN) segmentation of a 114×98×96 µm volume of zebra finch Area X brain tissue acquired with serial blockface EM at a voxel resolution of 9×9×20 nm [5]. Base FFN supervoxels (SVs) were agglomerated (Fig. 1a) via FFN resegmentation, with additional post-processing applied to the agglomeration graph to reduce merge and split errors [9]. We also used precomputed organelle probability maps for synaptic junctions and vesicle clouds [4] in some experiments.

**Fig. 1.**
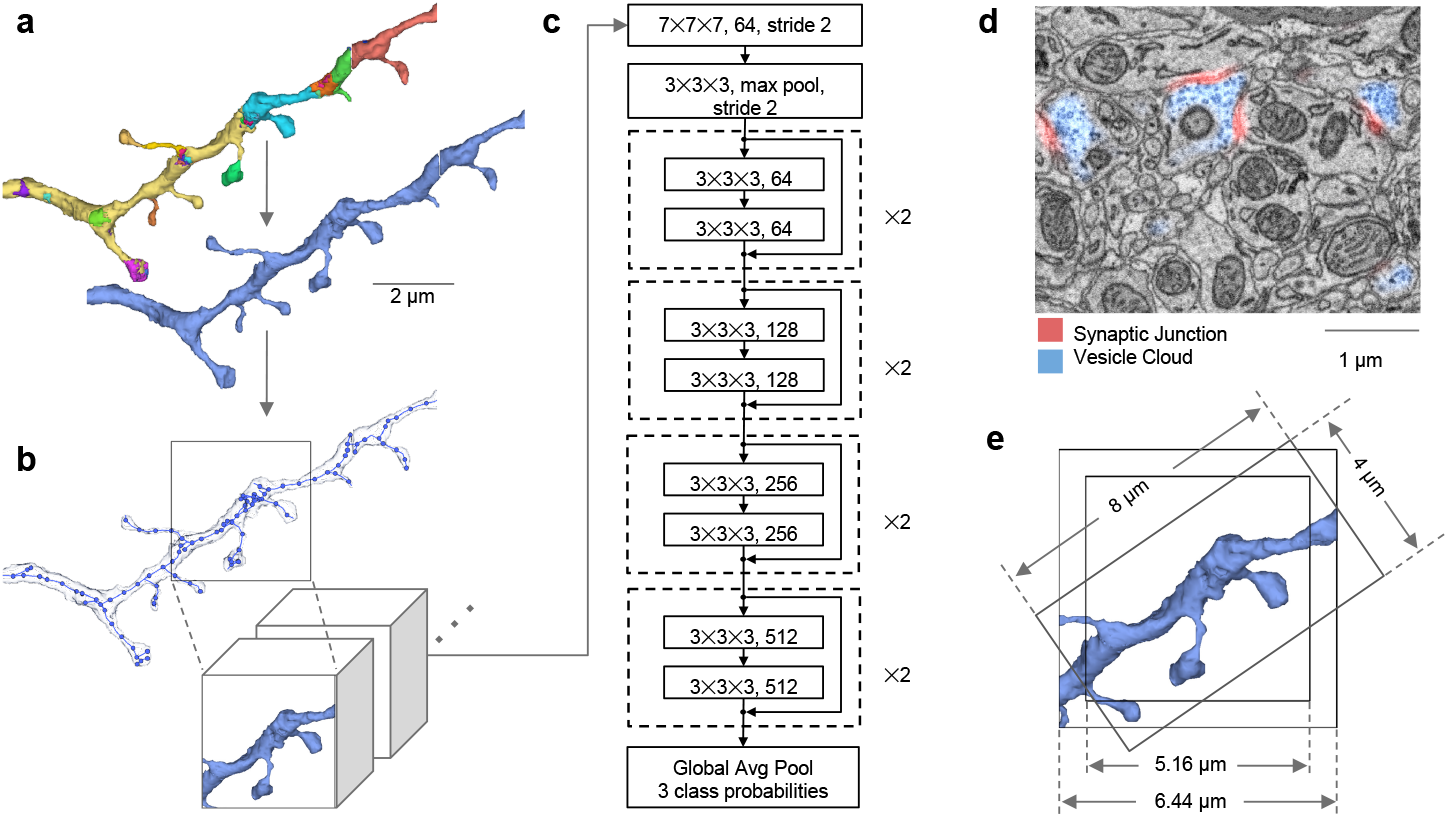
Neural subcompartment classification with 3d CNN. **(a)** The segmentation consists of base SVs (top, different colors) that were agglomerated into more complete neuron segments (bottom, solid) [5]. **(b)** Input FOVs are centered at node positions from automated skeletonization of the segmentation mask. **(c)** The classifier architecture is a 3d extension of a ResNet-18 CNN, and outputs probabilities for axon, dendrite, and soma subcompartment classes. **(d)** For some experiments, we provided additional input channels, e.g. the contrast normalized [20] EM image, or precomputed organelle probability maps [4]. **(e)** Illustration of the two primary FOV sizes used in our experiments, approximately 5.16 or 6.44 μm on a side. For comparison, we also illustrate the neurite-aligned 4×4×8 μm FOV employed by the previous CMN approach [12].

The agglomerated segmentation was skeletonized via TEASAR [19], and the resulting skeletons were sparsified to a mean inter-node spacing of 300 nm and eroded so that terminal nodes were at least 100 nm from the segment boundary. A subset of the objects in the volume were manually classified by human experts as axon, dendrite, or soma, of which 27 objects were used for training (32.8k axon, 8.4k dendrite, 7.5k soma nodes), 6 objects for validation (2.6k, 1.0k, 2.0k), and 28 objects for evaluation (29.3k, 38.2k, 32.7k).

Datasets are available from the CMN authors on request [12].

### 2.2 Classification of neural subcompartment with 3d CNNs

Classifier input fields of view (FOVs) were centered at neuron skeleton node locations, with the segment mask extracted from the neuron’s agglomerated segmentation (Fig. 1b). However, multiple axonal and dendritic processes from the same neuron sometimes pass close to each other, even if their connection point is far outside the FOV. Therefore, it was beneficial to remove segment mask components in the FOV that were not connected with the component at the center. Disconnected component removal was done at full 9×9×20 nm resolution, prior to downsampling the block to the network input resolution.

Classifier architectures were derived from the ResNet-18 CNN model [21], with convolution and pooling layers extended to 3d (Fig. 1c). Neuronal morphology was provided to classifiers as a 3d binary segment mask. When additional input channels (EM image, organelle masks; Fig. 1d) were provided, the segment mask was applied to the other channels instead of being provided separately, with areas outside the mask set to zero. Input data was provided at 36×36×40 nm resolution in blocks of 129 or 161 voxels on a side, for a total field of view of 4.64×4.64×5.16 or 5.80×5.80×6.44 μm respectively (Fig. 1e). Network output comprised probabilities for axon, dendrite, and soma sub-compartment classes.

For training, the input locations were class balanced by resampling skeleton nodes for underrepresented classes multiple times per epoch. We also applied random 3d rotations to the input as a training data augmentation. Networks were trained via stochastic gradient descent, with learning rate 0.003 and batch size 64 for 1.5M steps. For the best performing network, with two input channels and 6.44 μm field of view, the total number of trainable parameters was 33.2M.

### 2.3 Automated detection and correction of merge errors

We applied top-performing subcompartment predictions (Fig. 2) to further improve neuron reconstruction quality by detecting merge errors between different classes. FFN reconstruction biases base SVs (Fig. 1a) to have very few merge errors via an oversegmentation consensus procedure [5], so we focused on errors in SV agglomeration. Once agglomeration errors are identified and localized, they can be fixed efficiently by simply removing the bad agglomeration graph edges, either under human review [18] or automatically.

**Fig. 2.**
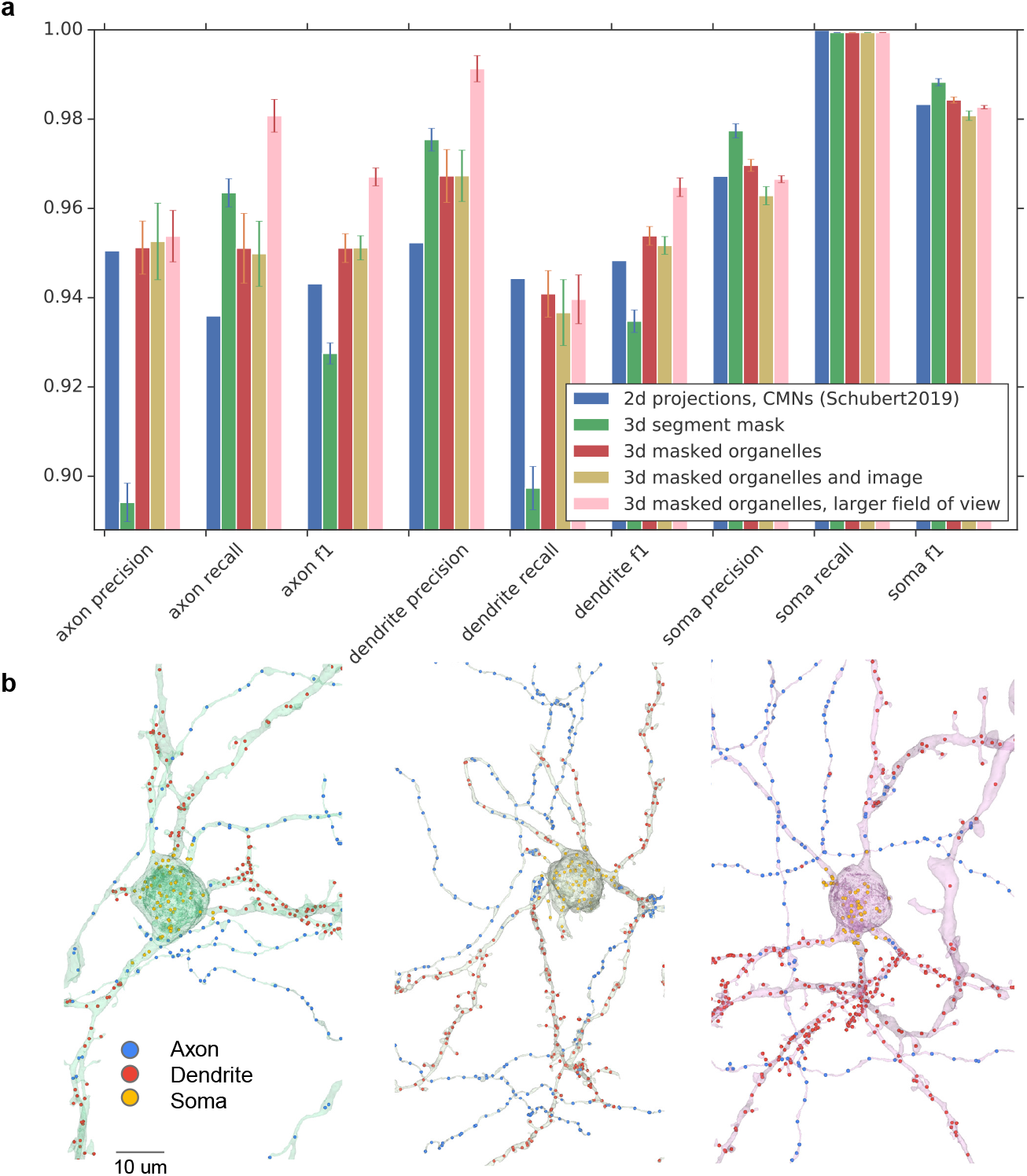
Subcompartment classification results. **(a)** Node classification performance on axon, dendrite, and soma labeled examples. **(b)** Skeleton node classifications of three automated neuron reconstructions outside the train, validation, and evaluation sets.

We used subcompartment predictions to identify all somas and branches, and then to detect and correct two classes of agglomeration errors: axon/dendrite branch merge errors, and soma/neurite merge errors. As ground truth, we manually identified 132 agglomerated neurons that contained merge errors and annotated their bad agglomeration graph edges. This yielded 473 branches, among which there were 83 branch merge errors and 56 soma merge errors. Together these represent a significant fraction of all merge errors identified through an exhaustive screening of the reconstruction.

### 2.4 Branch merge error correction by graph cut consistency score

Branch merge errors involve a mis-agglomerated axon and dendrite (Fig. 3a). Intuitively, branches that contain a merge error tend to have lower overall node prediction consistency (defined as weighted mean probability of dominant class type). Removing a bad agglomeration edge should improve the node consistency of the two resulting subgraphs.

**Fig. 3.**
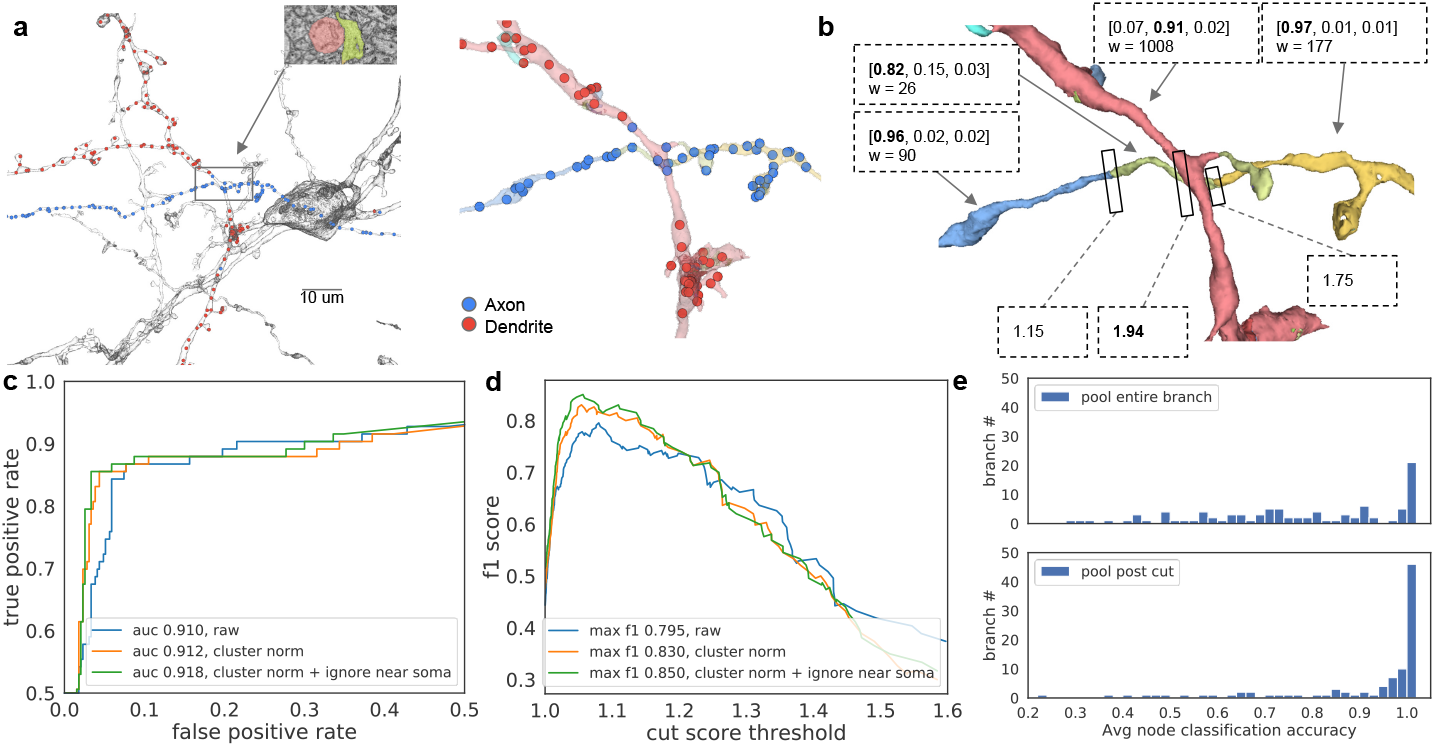
Correction of branch merge errors via subcompartment prediction. **(a)** Left, view of an agglomerated segment centered on a branch merge error, with the node predictions for the branch overlaid. Inset shows the EM image and overlaid base SVs for the merge. Right, zoomed in view of the merge error, with base SVs in different colors. **(b)** Node predictions are aggregated to get class probabilities [axon, dendrite, soma] and weight (node count) per SV. Three SVs are predicted axon, one dendrite, reflecting the merge error. Candidate neuron cuts are annotated with their consistency improvement scores (Eq. 1). **(c)** ROC plot showing detection performance as the cut score threshold is varied. Separate curves show three variants with different node reweighting (to address node clustering or nodes close to the soma). **(d)** The f1 of merge detection versus cut score threshold. **(e)** Branch-wise majority vote pooled class accuracy distribution before (top) and after (bottom) applying suggested cuts.

The input to our system is the skeleton of the agglomerated neuron with node class predictions, and the neuron’s agglomeration graph, where each skeleton node contains information about the base SV it belongs to. The workflow is as follows:

*Step 1:* Identify soma. If the segment has > 200 soma classified nodes, find the SV with the most soma nodes.

*Step 2:* Separate branches from soma. After removing the primary soma SV, each remaining subgraph of the agglomeration graph is considered a branch if it contains > 100 nodes.

*Step 3:* Compute node weights (optional). Automated skeletonization sometimes over-clusters nodes within thicker objects. Densely clustered nodes can optionally be down-weighted by 1 / (node count in 500 nm radius - 2) to discount the nodes in excess of the three expected within each 500 nm of clean path length. Furthermore, neurite nodes proximal to the soma (within 5-10 μm) tend to have inconsistent class predictions. These nodes can also be optionally down-weighted by 0.01, which effectively ignores them except in cases where soma proximal nodes are the only nodes on a branch.

*Step 4:* Group predictions. Node predictions are aggregated by base SV, to compute weighted mean class probabilities *PSV* and node count *w*_*SV*_ for each SV (Fig. 3b).

*Step 5:* Compute cut scores. Any cycles are first removed, then edges are traversed from leaf nodes in. At each edge, the branch is conceptually divided into subgraphs *G*_*leave*_ and *G*_*remain*_, and the “cut consistency score”, a measure of how many nodes belong to their respective majority classes post-versus pre-cut, is computed (Fig. 3b):

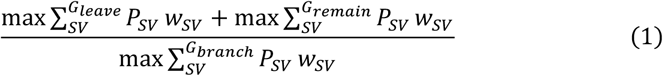

*Step 6:* Detection. The highest predicted cut score is thresholded to determine if the branch contains a merge error, with constraints that *G*_*leave*_ and *G*_*remain*_ must have different majority class types and their weighted sizes must be > 50.

*Step 7:* Correction (optional). The suggested agglomeration edge is removed, and majority vote pooling is performed within subcomponents. Branch pooled node prediction accuracy is compared pre- and post-cut (Fig. 3e).

### 2.5 Soma merge error correction by trajectory of primary neurite

The second error mode involves a neurite fragment that is mis-agglomerated with soma (Fig. 4). We observed that these errors can be fixed with a simple heuristic based on branch trajectory relative to the soma surface. The pipeline is as follows:

**Fig. 4.**
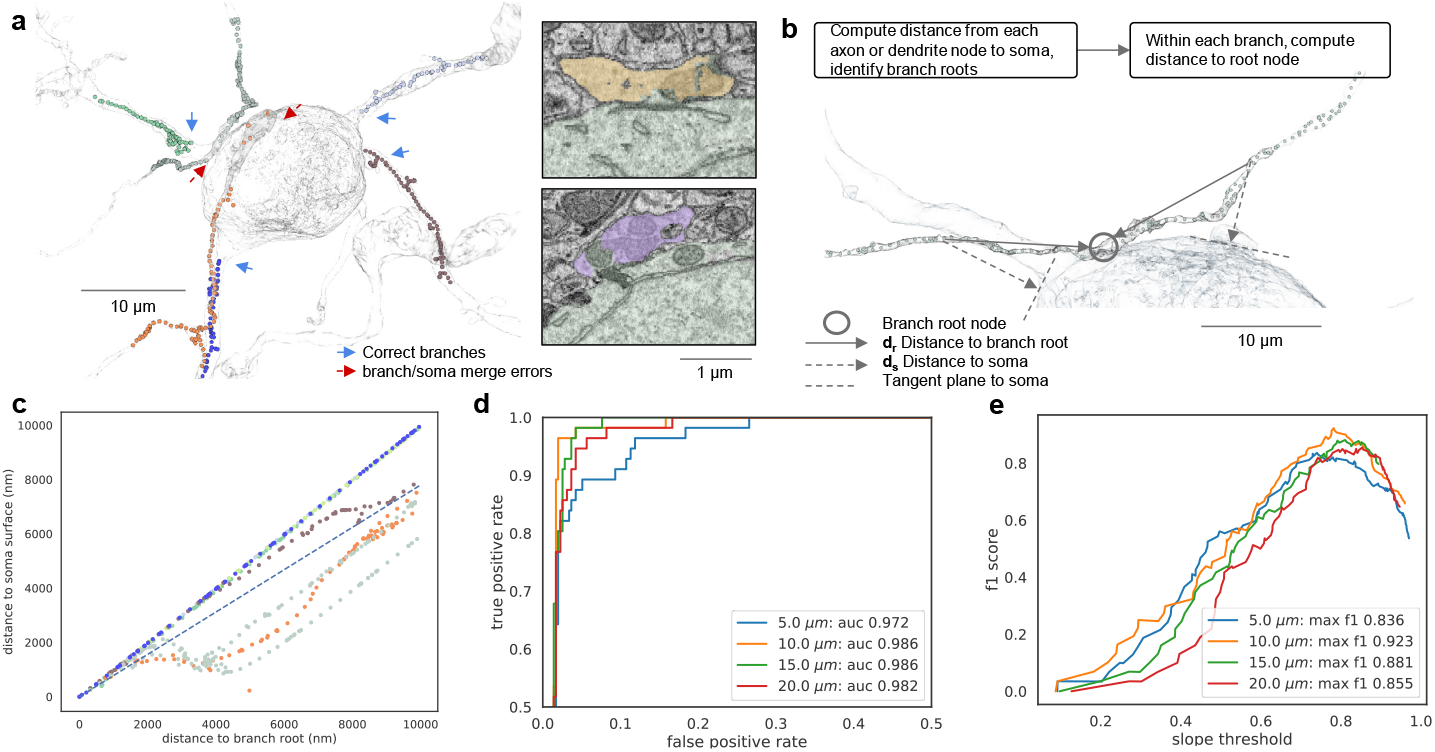
Correction of soma merge errors. **(a)** Example of a soma with multiple neurite branches, each in a different color. Two of the branches were erroneously agglomerated to the soma. **(b)** For each node along a branch, the distance to the nearest soma node is computed. The distance to the branch root (defined as the branch node closest to the soma) is also computed. **(c)** The soma distance versus branch root distance for the nodes comprising branches from (a), with matching color-coding. The dashed line of slope 0.78 separates the trajectories of correct branches that run primarily radially out from the soma, from the soma merge error branches that run primarily tangential. **(d)** ROC plot showing performance of merge error detection as slope threshold is varied. Separate curves show results with nodes at different distances from the soma included in the analysis. **(e)** The f1 of merge error detection versus slope threshold.

*Steps 1-2:* See branch merge detection pipeline above.

*Step 3:* Distance to soma. For each axon or dendrite node, compute the distance *d*_*s*_ to the nearest soma node.

*Step 4:* Distance to branch root. For each branch, the root is the node with minimum *d*_*s*_. Compute the distance *d*_*r*_ from each branch node to the root (Fig. 4b).

*Step 5:* Fit slope. For each branch, compute a linear fit to *d*_*s*_ versus *d*_*r*_ (Fig. 4c) for nodes within a tunable distance to soma. The slope of the fit is then thresholded to determine if a branch is a soma merge error (Fig. 4c-e).

## 3 Results

### 3.1 Subcompartment classification performance of 3d CNNs

We compared the performance of our 3d CNN classifiers to previous state-of-the-art results from CMNs [12], in terms of class-wise precision, recall, and f1 metrics (Fig. 2a). For each trained 3d CNN, we saved parameter checkpoints throughout the training period and screened them on a small manually labeled validation set. For most models, performance on the validation set approached or exceeded 0.99 on all metrics (not shown), but validation performance was useful for tracking convergence, confirming there was no overfitting to the training set, and for avoiding checkpoints where training temporarily became unstable. We then applied the ten checkpoints with highest validation accuracy to the larger evaluation set to compute the mean and standard deviation for each metric.

Compared with CMNs (Fig. 2a, blue), a network analyzing voxel representation of 3d segment shape alone was competitive (green). Adding vesicle cloud and synaptic junction organelle probability map channels allowed the 3d CNN to exceed state of the art (red). Interestingly, further adding the full EM image channel had negligible impact (yellow). Expanding the field of view for the masked organelles network from 5.16 μm to 6.44 μm yielded the best performing system tested (pink). We also tested expanding the FOV further, increasing the input resolution, increasing the CNN depth, and providing different input channel configurations; see supplemental Table S1.

Of the top ten checkpoints from the best performing model, the median overall node accuracy on the evaluation set was 97.1%, and mean f1 across classes was 0.972. We then used this median checkpoint to predict node classes for reconstructed neurons and fragments throughout the entire volume. Predicted skeletons demonstrate good class consistency within soma and neurites, with some ambiguity at the interface between branch and soma (Fig. 2b). Based on the predictions, the volume contains 3.25 m total axon path length, and 0.79 m dendrite path length, a ratio of 4:1 that is similar to the 5:1 ratio previously reported [12]. However, the total path length here significantly exceeds that previously reported, probably due to differences in skeleton sparsity, so the absolute lengths here should be considered an upper bound.

### 3.2 Agglomeration merge error detection and correction

We fed subcompartment predictions back to detect and correct two classes of reconstruction merge errors that occur during SV agglomeration: axon/dendrite branch merges, and soma/neurite merges.

The branch merge error correction system is based on analyzing the predicted subcompartment class consistency of agglomerated segments, with and without candidate cuts applied (Fig. 3a-b). We first considered branch merge error detection performance, and plotted the receiver operating characteristic (ROC) curve by varying the cut score threshold (Fig. 3c-d). In areas with many small SVs, several nearby cut candidates can have equivalent impact, so predicted cuts that fell within four agglomeration graph edges of ground truth cuts were considered correctly detected. The best detection performance was at 1.05 cut score threshold, with f1 of 0.850 (see also Table S2).

Merge error detections can be used to flag the location for human review. We also calculated the node prediction accuracy improvement after directly applying the suggested cut. For the 96 branches with either a predicted merge or ground truth merge, we manually determined their nodewise ground truth class as axon or dendrite, then performed majority vote predicted class pooling before and after applying predicted cuts. Comparing class pooled accuracy pre- and post-cut (Fig 3e), the mean node prediction accuracy improves from 0.804 to 0.886.

We addressed the second category of agglomeration merge errors, between somas and nearby neurites, by analyzing the trajectory of the branch relative to the somal surface (Fig. 4a-c). We found the best performance is achieved by sampling skeleton nodes within the initial 10 μm from the soma, yielding an f1 of 0.923 at a slope of 0.78. (Fig. 4d-e; see also Table S3).

Combined, branch and soma merge analyses detected 90.6% of merge errors, with a false positive rate of only 2.7%.

## 4 Conclusions & Discussion

To make volume EM datasets of brain tissue easy to analyze at scale, it is crucial to reduce the data they contain to more compact and semantically meaningful representations. Segmentation and synapse detection provide an important first step in this process. Here we presented a system that can provide further information about the biological identity of neurites by predicting subcompartment types, and feed back to the preceding reconstruction stage through automated correction of agglomeration errors.

We expect this approach to be useful for brain circuit analyses, and to be applicable to diverse datasets. We also anticipate that the approach could be extended to finer grained subcompartment classification. For example, the subcompartment localization of a postsynaptic site on e.g. a dendritic spine, dendritic shaft, soma, or axon initial segment is linked to both the synapse’s functional impact as well as the identity of its presynaptic partner [22, 23]. Another related application is in the identification of neuronal subtypes, whose shared structural and functional properties can enhance connectome interpretability by organizing thousands of individual neurons into a reduced complement of conceptual roles [24–26].

The primary advantages of our system are its simplicity, and its ability to capture complete local information about a neurite, resulting in a new state of the art. A fundamental limitation is that processing efficiency drops with increasing field of view as the neurite of interest fills a progressively smaller fraction of the voxels that need to be processed. This limitation could be mitigated by using an alternative representation of sparse 3d data [27–31].

## Acknowledgments

We thank Philipp Schubert and Jörgen Kornfeld for sharing detailed CMN results and training data, as well as the EM volume. We thank Jeremy Maitin-Shepard and the anonymous reviewers for comments on the manuscript.

## Supplemental Material

### Node classification ablations and extensions

Table S1 presents performance metrics on the evaluation set for 15 node classification experiments, as well as the performance of the previous state-of-the-art CMNs [12]. Bold rows indicate data already plotted in the main text (Fig. 2).

Rows 5-7 and 9 focus on changing the input channels. Rows 5 and 6 show the relative performance of using only synaptic junction or vesicle cloud probability maps alone, rather than both together (row 3); the vesicle clouds perform better, but neither is sufficient on its own. Rows 7 and 9 show that adding mitochondria probability maps does not improve performance, and may cause some degradation.

Row 10 shows that further expanding the field of view (FOV) to 193 voxels on a side degrades performance relative to the 161 voxel model (row 8). As FOV increases, the neuron segment mask is increasingly sparse, i.e. the input is increasingly empty. This may cause the model to train less smoothly, consistent with the larger checkpoint variance.

Rows 11 and 12 show that increasing input resolution to 18×18×20 nm degrades performance relative to the 36×36×40 nm models (e.g. row 3). The number of voxels in the input block for the higher resolution models remained the same, so the resulting reduction in FOV in terms of microns likely explains the degradation. Interestingly, adding the full EM image does improve the performance of these higher resolution models, in contrast to the lower resolution models where adding the image has little effect (rows 3 and 4). This is consistent with our observation that as human viewers we find a significant amount of detail is lost in the EM image when downsampling from 18×18×20 to 36×36×40 nm.

Rows 13-16 repeat several of the experiments using a deeper ResNet-50 network architecture. Overall, the results are similar to ResNet-18 performance. The best performing ResNet-18 model performs somewhat better than its Res-Net-50 equivalent (rows 8 and 16) and is significantly less computationally intensive, so we favored ResNet-18 in experiments and production.

**Table S1.**
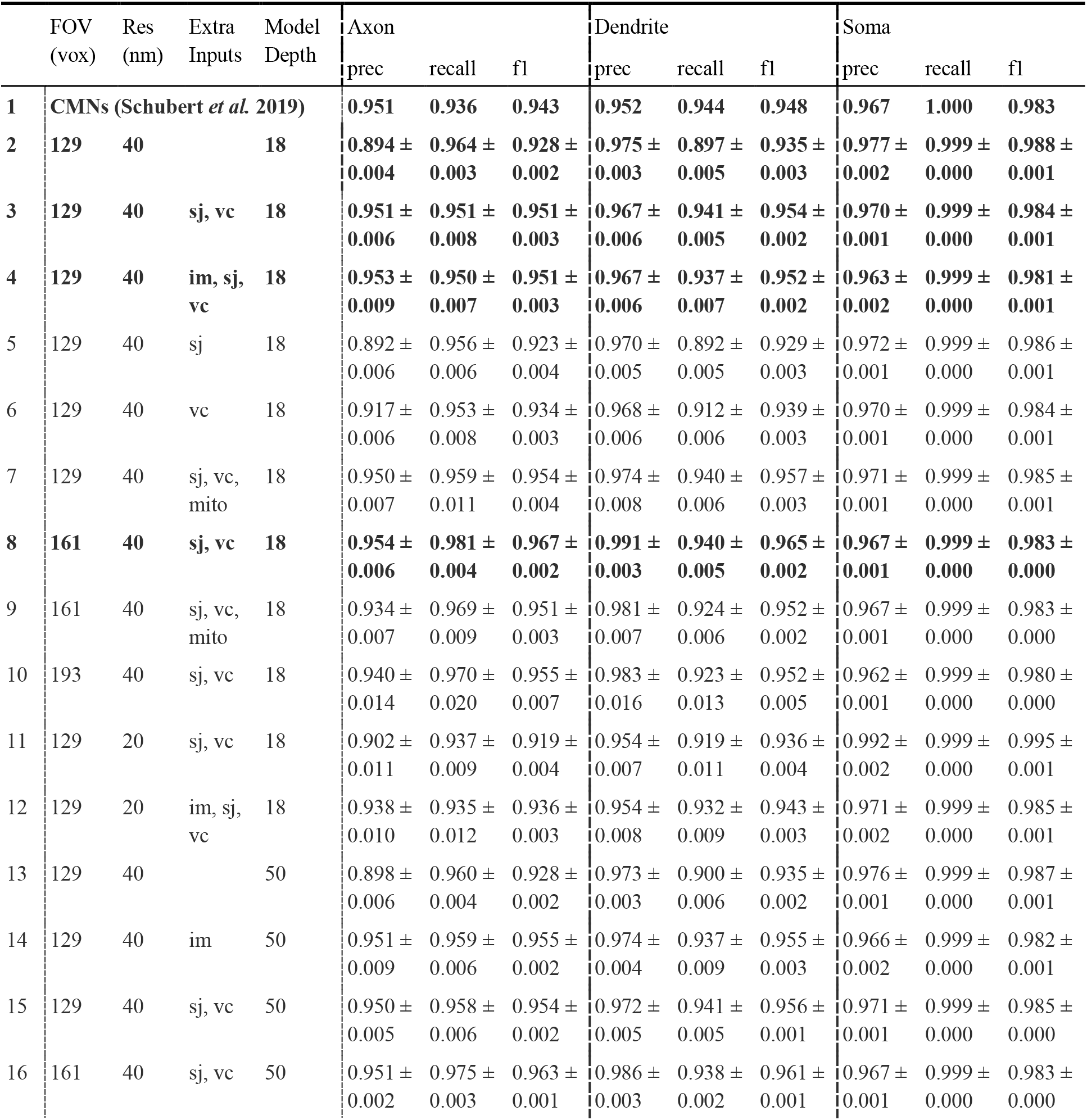
Node classification ablations and extensions vox: voxels on a side; Res: roughly isotropic resolution; prec: precision; im: EM image; sj: synaptic junctions; vc: vesicle clouds; mito: mitochondria

### Detailed branch merge detection metrics

Table S2 shows the detailed performance metrics for the best performing (in terms of f1 metric) cut score threshold settings of the branch merge detection pipeline, corresponding to data plotted in the main text (Fig. 3c-d).

**Table S2.** Branch merge detection performance (best f1)

| Node reweighting | Precision | Recall | F1 | AUC |
| --- | --- | --- | --- | --- |
| no reweighting | 0.753 | 0.843 | 0.795 | 0.910 |
| clusters | 0.807 | 0.855 | 0.830 | 0.912 |
| clusters, near soma | 0.845 | 0.855 | **0.850** | 0.918 |

### Detailed soma merge detection metrics

Table S3 shows the detailed performance metrics for the best performing (in terms of f1 metric) slope threshold settings of the soma merge detection pipeline, corresponding to data plotted in the main text (Fig. 4d-e). After excluding predicted soma merge error branches, we were able to identify a single correct axonal branch 82.4% of the time.

**Table S3.**
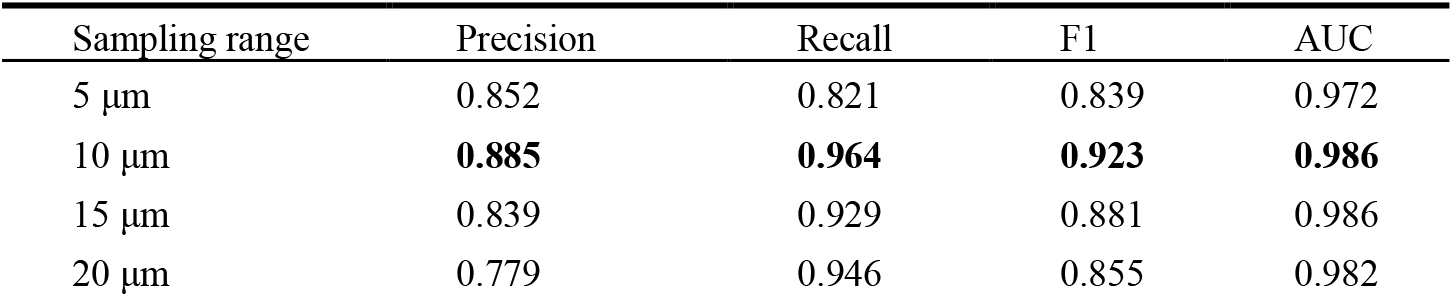
Soma merge detection performance (best f1)

## References

1. Zheng Z, Lauritzen JS, Perlman E, et al (2018) A Complete Electron Microscopy Volume of the Brain of Adult Drosophila melanogaster. Cell 174:730–743.e22

2. Dorkenwald S, Turner NL, Macrina T, et al (2019) Binary and analog variation of synapses between cortical pyramidal neurons. bioRxiv 2019.12.29.890319

3. Shan Xu C, Januszewski M, Lu Z, et al (2020) A Connectome of the Adult Drosophila Central Brain. bioRxiv 2020.01.21.911859

4. Dorkenwald S, Schubert PJ, Killinger MF, et al (2017) Automated synaptic connectivity inference for volume electron microscopy. Nat Methods 14:435–442

5. Januszewski M, Kornfeld J, Li PH, et al (2018) High-precision automated reconstruction of neurons with flood-filling networks. Nat Methods 15:605–610

6. Li PH, Lindsey LF, Januszewski M, et al (2019) Automated Reconstruction of a Serial-Section EM Drosophila Brain with Flood-Filling Networks and Local Realignment. bioRxiv 605634

7. Buhmann J, Sheridan A, Gerhard S, et al (2019) Automatic Detection of Synaptic Partners in a Whole-Brain Drosophila EM Dataset. bioRxiv 2019.12.12.874172

8. Dasgupta S, Stevens CF, Navlakha S (2017) A neural algorithm for a fundamental computing problem. Science 358:793–796

9. Kornfeld JM, Januszewski M, Schubert PJ, et al (2020) An anatomical substrate of credit assignment in reinforcement learning. bioRxiv 2020.02.18.954354

10. Swanson LW, Lichtman JW (2016) From Cajal to Connectome and Beyond. Annu Rev Neurosci 39:197–216

11. Motta A, Berning M, Boergens KM, et al (2019) Dense connectomic reconstruction in layer 4 of the somatosensory cortex. Science, 366(6469).

12. Schubert PJ, Dorkenwald S, Januszewski M, et al (2019) Learning cellular morphology with neural networks. Nat Commun 10:2736

13. Meirovitch Y, Matveev A, Saribekyan H, et al (2016) A Multi-Pass Approach to Large-Scale Connectomics. arXiv [q-bio.QM]

14. Rolnick D, Meirovitch Y, Parag T, et al (2017) Morphological Error Detection in 3D Segmentations. arXiv [cs.CV]

15. Haehn D, Kaynig V, Tompkin J (2018) Guided proofreading of automatic segmentations for connectomics. Proceedings of the IEEE Conference on Computer Vision and Pattern Recognition

16. Krasowski NE, Beier T, et al (2017). Neuron segmentation with high-level biological priors. IEEE transactions on medical imaging, 37(4), 829–839.

17. Pape C, Matskevych A, et al (2019). Leveraging domain knowledge to improve microscopy image segmentation with lifted multicuts. Frontiers in Computer Science, 1, 6.

18. Hubbard PM, Berg S, Zhao T, et al (2020) Accelerated EM connectome reconstruction using 3D visualization and segmentation graphs. BioRxiv

19. Sato M, Bitter I, Bender MA, et al (2000) TEASAR: tree-structure extraction algorithm for accurate and robust skeletons. In: Proceedings the Eighth Pacific Conference on Computer Graphics and Applications. pp 281–449

20. Zuiderveld K (1994) Contrast Limited Adaptive Histogram Equalization. In: Heckbert PS (ed) Graphics Gems IV. Academic Press Professional, Inc., San Diego, CA, USA, pp 474–485

21. He K, Zhang X, Ren S, Sun J (2016) Deep Residual Learning for Image Recognition. In: Proceedings of the IEEE Conference on Computer Vision and Pattern Recognition. pp 770–778

22. Petilla Interneuron Nomenclature Group, Ascoli GA, Alonso-Nanclares L, et al (2008) Petilla terminology: nomenclature of features of GABAergic interneurons of the cerebral cortex. Nat Rev Neurosci 9:557–568

23. Contreras A, Hines DJ, Hines RM (2019) Molecular Specialization of GABAergic Synapses on the Soma and Axon in Cortical and Hippocampal Circuit Function and Dysfunction. Front Mol Neurosci 12:154

24. Jiang X, Shen S, Cadwell CR, et al (2015) Principles of connectivity among morphologically defined cell types in adult neocortex. Science 350:aac9462

25. Gouwens NW, Sorensen SA, Baftizadeh F, et al (2020) Toward an integrated classification of neuronal cell types: morphoelectric and transcriptomic characterization of individual GABAergic cortical neurons. bioRxiv 2020.02.03.932244

26. Grünert U, Martin PR (2020) Cell types and cell circuits in human and non-human primate retina. Prog Retin Eye Res 100844

27. Kipf TN, Welling M (2016) Semi-Supervised Classification with Graph Convolutional Networks. arXiv [cs.LG]

28. Qi CR, Su H, Mo K, Guibas LJ (2016) PointNet: Deep Learning on Point Sets for 3D Classification and Segmentation. arXiv [cs.CV]

29. Riegler G, Osman Ulusoy A (2017) Octnet: Learning deep 3d representations at high resolutions. Proceedings of the IEEE Conference on Computer Vision and Pattern Recognition

30. Mescheder L, Oechsle M, Niemeyer M (2019) Occupancy networks: Learning 3d reconstruction in function space. Proceedings of the IEEE Conference on Computer Vision and Pattern Recognition

31. Graham B, Engelcke M, & Van Der Maaten L (2018). 3d semantic segmentation with submanifold sparse convolutional networks. In Proceedings of the IEEE conference on computer vision and pattern recognition (pp. 9224–9232).

